# Renal adaptation to primary aldosteronism requires GDF15 to control plasma potassium level in mice and human

**DOI:** 10.64898/2026.10.18.759915

**Authors:** Justine Billiet, Luciana Morla, Emmanuelle Vidal-Petiot, Emmanuel Wirth, Maude Laney, Stéphanie Baron, Laurence Amar, Lydie Cheval, Gilles Crambert

## Abstract

**Background:** Primary aldosteronism (PA), the most common secondary cause of hypertension, results from autonomous aldosterone production, driving renal Na□ retention and K□ secretion. Paradoxically, only 9–37% of PA patients develop hypokalemia, suggesting adaptive mechanisms. Growth differentiation factor 15 (GDF15) promotes intercalated cell type A (ICA) proliferation and K□ reabsorption in the collecting duct, but its role in PA is unexplored.

**Methods:** We measured urinary and plasma GDF15 in 81 PA patients (two centers) and 41 controls. A murine model of chronic mineralocorticoid excess (DOCP treatment) was used in wild-type (WT) and GDF15 knockout (GDF15KO) mice to assess renal adaptation.

**Results:** PA patients exhibited 3-fold higher urinary GDF15 and 1.6-fold higher plasma GDF15 than controls, with normalization post-treatment. 24-hour urinary GDF15 correlated with urinary aldosterone (ρ=0.35, p<0.01), age, and BMI (Body Mass Index), but not plasma K□. WT mice developed hypokalemia following DOCP administration (plasma K□ 3.5±0.1 mM vs. 4.3±0.1 mM in controls, p<0.01). In GDF15KO mice DOCP-induced hypokalemia was more severe, indicating a protective role of GDF15. In WT mice, DOCP increased 24-hour urinary GDF15 (2-fold) and GDF15 expression in the collecting duct. Transcriptomic analysis revealed an inverted transcriptional response to chronic mineralocorticoid in ICA and PC according to the genotype. This could be related to the increase of ICA number in WT, an effect abolished in GDF15KO mice.

**Conclusions:** These results indicate that GDF15 is involved in the renal response to mineralocorticoid excess, likely through stimulation of ICA proliferation, which may promote potassium reabsorption in the collecting duct and help mitigate aldosterone-mediated K□ loss.

## Introduction

Primary aldosteronism (PA) is the most common cause of secondary hypertension with a prevalence of 5-10%^1^ in hypertensive patients and 20%^2,3^ in patients with resistant hypertension. It is associated with more severe target organ damage compared with essential hypertension. PA is characterized by autonomous aldosterone overproduction by the zona glomerulosa of the cortex of adrenal gland which can be either unilateral (aldosterone producing adenoma or Conn’s syndrome) or bilateral (bilateral adrenal hyperplasia). Familial hyperaldosteronism causes by variants with autosomal dominant inheritance (types I to IV)^4^ also exist. In normal condition, this steroid hormone is produced in response to salt loss, hypovolemia, hyperkalemia or acidosis (for review see^5^) to regulates blood pressure, plasma K^+^ level^6^ or acid/base balance^7^ whereas in PA, aldosterone is overproduced independently of its physiological triggers. Aldosterone mainly regulates electrolyte homeostasis through genomic effects due to activation of the mineralocorticoid receptor (MR) in the kidney but non-genomic effects have also been reported^8,9^.

In kidneys, MR is expressed by the principal cells (PC) of the collecting duct (CD) which are responsible for the reabsorption of sodium (Na^+^), water and the secretion of potassium (K^+^). It is also found in the intercalated cells type A (ICA)^10^ where its activation stimulates H^+^ secretion. Upon activation of the MR, the activity and expression of the Epithelial Na Channel (ENaC) are strongly increased at the apical membrane of the PC as are those of the Na,K-ATPase at its basolateral side. MR activation in ICA increases the number of H^+^-pump at their apical side^10^. In cases of aldosterone excess, the reabsorption of Na^+^ is increased. This leads to a brief rise in plasma osmolarity which triggers the production of vasopressin and stimulates water reabsorption through aquaporin 2 (AQP2), overall leading to expand extracellular volume, and elevated blood pressure. The increased reabsorption of Na^+^ by ENaC in PC also depolarizes the apical membrane, which favors K^+^ secretion by the channel Renal Outer Medullary potassium channel (ROMK) and may promote hypokalemia. If PA is responsible for hypertension in all patients, only a minority of cases (around 10 to 40% depending on the studies) are hypokalemic^11,12^. Hypokalemia remains a marker of disease severity, being more common in aldosterone-producing adenomas (52%) than in bilateral adrenal hyperplasia (22%)^1^ and is correlated with a higher risk of developing cardiovascular issues^12^. The development of overt hypokalemia may be mitigated by compensatory mechanisms counteracting the kaliuretic effect of aldosterone.

Potassium homeostasis relies on two complementary systems^13^. The first is internal balance, which involves transcellular distribution of K^+^. This rapid regulation process avoids hyperkalemia after each meal by temporary storage of K^+^, primarily in muscle cells with progressive release into the extracellular compartment thereafter. The second, longer term regulation system, is external balance, mediated by the kidneys and gut, which regulate the intake and excretion of K^+^ from the body. Kidneys can limit hypokalemia by reducing K^+^ excretion through different mechanisms. Hypokalemia induced by a low K^+^ diet has been shown to induce the production of ROS compounds that inhibit ROMK^14^. Hypokalemia has also been shown to activate Na^+^ reabsorption, upstream of the collecting duct, by the Na/Cl-cotransporter (NCC)^15^ in the distal convoluted tubule. Consequently, less Na^+^ is delivered to the collecting duct and ENaC is less active which reduces the secretion of K^+^. Another mechanism involves the stimulation of the K^+^ reabsorption by the H,K-ATPase type 2 (HKA2) expressed in ICA^16^. We have demonstrated that under low K^+^ diet, leading to hypokalemia, a growth factor, GDF15, was overexpressed by the different segments of the nephron^17^. It has also been shown that in the collecting duct, GDF15 is produced by PC^18^. This factor is responsible for the increased expression of the HKA2 and proliferation of the ICA^19^, leading to a higher ability for K^+^ reabsorption. While these adaptive mechanisms have been described, the role of GDF15 in counteracting aldosterone-induced kaliuresis in PA remains unexplored. Here, we hypothesized that GDF15-mediated ICA proliferation may be stimulated in response to chronic aldosterone excess, as a renal adaptative mechanism to protect against hypokalemia. To test this hypothesis, we first measured GDF15 levels in PA patients and then investigated its role in a murine model of chronic mineralocorticoid excess.

## Material and methods

### Patients

This ambispective study was conducted using patient data and samples collected at two different centers: the Department of Physiology and Hypertension center at Bichat hospital (center 1) and at the Hypertension Research and Care Unit at Georges Pompidou European hospital (center 2). Eligible participants were aged > 18 years-old at inclusion and signed a written informed consent for data collection and scientific use of their anonymized data. The study was performed according to the Declaration of Helsinki and was approved by the local ethics committee (IRB00006477, subject N° CER 2025-314). The study included a PA group (n=42 from center 1 and n=39 from center 2) with confirmed diagnosis of primary aldosteronism, a group cured after surgical treatment of PA (n=13 from center 1) and a control group composed of healthy kidney donor candidates (n=41 from center 1). All patients underwent standardized plasma and 24-hour urine measurements after withdrawal of interfering treatment for at least two weeks to ascertain PA diagnosis^20^. To focus on the most severe cases, we initially selected PA patients with hypokalemia (plasma K^+^ values <3.5 mM, center 1) but later also included less severe cases with normokalemic PA patients from center 2 (n=13 of the 39).

In order to take into account potential inaccuracy of 24-hour urine collections, urinary creatinine excretion was calculated, and urine parameters were corrected using the mean expected daily creatinine excretion (0.15 mmol/kg in women and 0.19 mmol/kg in men as reported by Forni et al)^21^.

GDF15 were measured by ELISA test (DGD150, R&D Systems) in 24-hour urine and fasting plasma samples.

### Data and statistical analysis

Results are presented as medians with interquartile ranges (IQR) for skewed variables, and percentages for categorical variables.

The normality of the variable distribution was assessed using Shapiro-Wilk test. Noteworthy, 24-hour urinary aldosterone and 24-hour urinary and plasma GDF15 levels did not follow a normal distribution neither before nor after logarithmic transformation. Comparisons between two groups were performed using a Student T-test (normal distribution, Shapiro-Wilk test p<0.05) or its non-parametric equivalent, the Mann-Whitney test (non-normal distribution, Shapiro-Wilk test p>0.05). Comparison involving more than two groups were conducted using one-way ANOVA (normal distribution, Shapiro-Wilk test p<0.05) or its non-parametric equivalent, the Kruskal-Wallis test (non-normal distribution, Shapiro-Wilk test p>0.05). Proportions were compared using the χ2 test. When the differences between groups were significant, we measured the size effect (Cohen’s d).

Because 24-hour urinary GDF15 levels did not follow a normal distribution, Spearman rank correlation analyses were performed to examine the associations of GDF15 levels with the explanatory variables (sex, age, BMI, plasma K^+^, 24-hour urinary K^+^ and 24-hour urinary aldosterone). Spearman’s rho [95% CI] was estimated via bias-corrected and accelerated (BCa) bootstrap resampling (2,000 replications). Patients with missing data were excluded from this analysis.

For the multiple linear regression (conducted in RStudio), explanatory variables were selected based on their clinical and pathophysiological relevance. The analysis excluded patients with missing data for one of the parameters. Collinearity among the explanatory variables was assessed using the variance inflation factor (VIF). All VIF values remained close to 1, indicating no significant collinearity. Model quality was evaluated using the adjusted R^2^. For all analyses, differences with p<0.05 were considered significant.

### Animals and treatment

Experiments were performed on male C57BL/6J wild type (WT) and knock out mice for Gdf15 gene (GDF15KO). The GDF15KO mice were first provided by Dr. Se-Jin Lee (John Hopkins University, Baltimore, MD) and backcrossed with C57BL/6J^22^. They are regularly backcrossed with wild-type C57BL/6J male mice from Janvier Labs to avoid genetic deviation. The animals are kept at CEF (Centre d’Explorations Fonctionnelles of the Cordeliers Research Center, Agreement no. A75-06-12).

Animals were fed a standard laboratory diet (0.3% of Na^+^ and 0.6% of K^+^; UPAE, INRA, Jouy-en-Jossas, France). They were treated with a single intramuscular injection of deoxycorticosterone pivalate (DOCP) an analog of aldosterone (75µg/g of body weight) in both thighs to respect volume administration^23,24^ and analyzed at 2, 9, 16, 21 and 28 days post-injection. To test the effect of a K^+^ load, we fed mice with a high potassium diet (5% of K^+^) either 7 days before DOCP-treatment and 2 days after, or 2 days after DOCP-treatment for 7 days.

### TD-NMR

The injection was validated with the measure of the free fluid with the time-domain nuclear magnetic resonance (TD-NMR) (Minispec, LF50, Bruker)^23^. All mice were habituated to TD-NMR analysis before the measurements. Mice were weighted after emptying their bladder before the measurements.

### Metabolic analysis

To record physiological parameters, mice were placed in metabolic cages (Techniplast, France) for 2 days of adaptation and were fed a standard laboratory diet (0.3 % Na ^+^ and 0.6 % K^+^; UPAE, INRA, Jouy-en-Jossas, France) or a high potassium diet (5% of K^+^). Plasma electrolytes were measured by retro-orbital puncture on anesthetized animal (a mix of xylazine and ketamine, respectively 10mg/kg and 100mg/kg) with an Epoc pH/blood-gas analyzer (Siemens Health-ineers).

Urinary creatinine concentrations were determined using an automatic analyzer (Indiko plus, Thermoscientific). Urinary K^+^ and Na^+^ concentrations were determined by flame photometry (M420, Sherwood Scientific, France). Urine GDF15 were measured by ELISA test (MGD150, R&D Systems).

### Aldosterone measurement in plasma

Plasma samples of mice were collected in control condition and at different time post treatment. Aldosterone was measured by mass spectrophotometry (Physiology Department, Hôpital Européen Georges Pompidou, Paris, France).

### Muscle potassium content

Gastrocnemius were removed to measure their potassium content. After muscles grinding in 1ml of water, the extracts were incubated overnight with trichloroacetic acid 20% (TCA) before centrifugation (12000G, 10 minutes at 4°C). The supernatants were then measured by flame photometry (M420, Sherwood Scientific, France).

### RNA sequencing, mRNA and protein expression

RNAs were extracted from whole kidneys of WT and GDF15KO mice either under control condition or 16 days post-treatment with DOCP using the TRI reagent (Invitrogen, France) following the manufacturer’s instructions. The quality of the extracted RNAs was checked using Agilent TapeStation 4200 (Agilent Technologies) and considered of good quality for a RIN > 7. The samples were then sent to the facility iGenSeq of the Institut du Cerveau et de la Moelle Epinière (Hôpital de la Pitié Salpêtrière, Paris, France) and processed for RNA sequencing using the Stranded mRNA kit (Illumina). The results have been deposited on the GEO website of the NCBI (GSE348216).

### Quantitative PCR

Segments of the nephron were microdissected from liberase-treated kidneys according to morphological parameters under binocular loupes. RNA extraction was performed using RNeasy micro kit (Qiagen). RNAs of kidney were extracted by TRI reagent (Invitrogen) following manufacturer’s instructions. Then, 1µg of RNA were retro-transcribed into cDNA according to manufacturer’s instructions (Roche Diagnosis). Real time PCRs were performed on a LightCycler (Roche Diagnostics). The expression of the house keeping gene Rpl26 is used to normalize the results. In each run, a standard curve was obtained using serial dilution of stock cDNA.

### Immunolabeling on isolated OMCD

Twenty outer medullary collecting ducts were isolated from liberase–treated kidney and transferred on Super Frost Plus glass slides and treated as described recently^18^. Slides were incubated overnight at 4°C with the primary antibody (anti-AE1 1/500 a gift from CA Wagner or catalog. no. 20112; Cell Signaling). Slides were mounted with Vectashield and observed by confocal microscopy (Zeiss observer. ZI, LSM710). For cell and doublet counting, 3D reconstructions from all stacks were performed (Image J) from images of AE1-labelled OMCDs acquired by confocal microscopy^18^. ICA were identified as AE1+ cells using anti-AE1 antibody. The total number of cells was also counted via DAPI staining (1/1000, D9542-1MG, Sigma). Seven or eight tubules per animal were analyzed.

### Data and statistical analysis

The number of mice used in these experiments are indicated in the legends (n). Results are presented as mean±sem or median with interquartile range according to the normality of the values. The normality of the result distribution was tested using Shapiro-Wilk test. Comparisons between two groups were performed using a Student T-test (paired or unpaired) or its non-parametric equivalent, the Mann-Whitney test depending on the normality of the distribution. Comparison involving more than two groups were conducted using one or two-way ANOVA. For all analyses, differences with p<0.05 were considered significant.

## Results

### GDF15 level is increased in PA patients

To investigate the level of GDF15 in PA patients compared to cured patients and to healthy kidney donor candidates (control group), we obtained clinical data, urine and plasma samples from center 1. The general characteristics of the different groups are presented in Table 1. Noteworthy, plasma K^+^ values are lower in PA patients compare to both the control and the PA post-treatment groups (3.03±0.04 mmol/l vs 3.79±0.04 mmol/l and 3.88±0.05 mmol/l, respectively) and 24-hour urinary K^+^ excretion is 1.6-fold higher in both control groups. Plasma renin and urinary aldosterone measurements were not performed in the kidney donor control group. As expected, in PA patients, plasma renin is 3-4 times lower than in the PA post-treatment group and their 24-hour urinary aldosterone excretion is 5.9-fold increased. Interestingly, 24-hour urinary excretion and plasma level of GDF15 are 3-fold (p<0.05, Cohen’s d=-1.38) and 1.6-fold increased (p<0.01, Cohen’s d=-0.83), respectively in PA patients compared to the control group (Figure 1A and B). In PA post-treatment group, 24-hour urinary excretion and plasma level of GDF15 are similar to the control group (Figure 1A and B). In three patients, values were available before and after treatment. As shown in Figure 1C, as expected, their plasma K^+^ levels return to the physiological values (3,5<plasma K^+^<5 mmol/L) after treatment and their 24-hour urinary excretion of GDF15 decreased to reach similar values than those before the treatment (Figure 1D).

**Figure 1.**
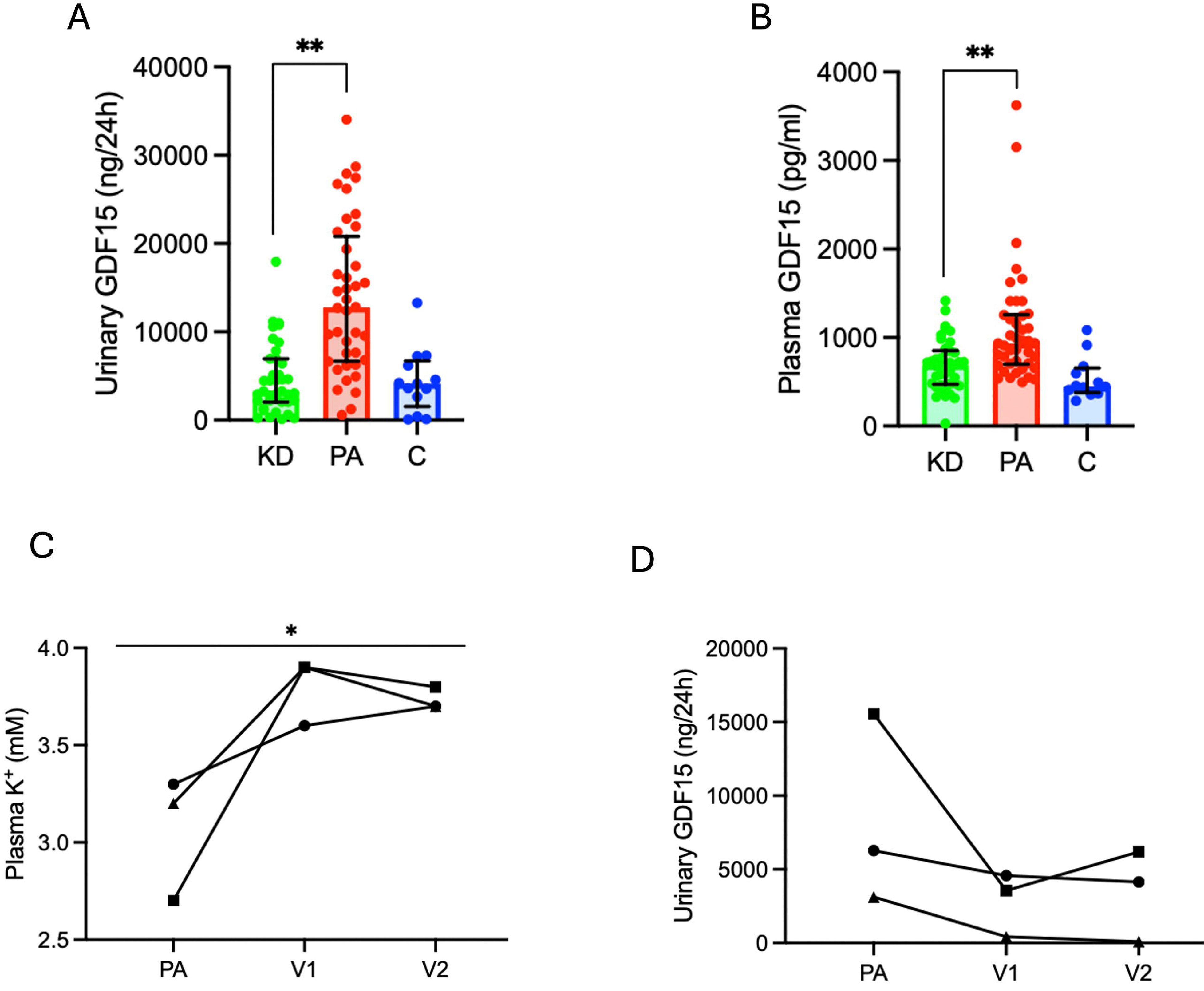
GDF15 is enhanced in patients with primary aldosteronism. A) Total excretion of GDF15 and B) plasma GDF15 in kidney donor candidates (KD), patients with primary aldosteronism (PA) and patients cured after treatment (C) from center 1. Results are shown as median with interquartile and analyzed by a Kruskal-Wallis test and followed by a Dunn’s multiple comparisons test with KD group as control. **p.value<0.01. C) Plasma K^+^ and D) total excretion of GDF15 of three patients before and after treatment with V1 and V2: first and second visit post-surgery. Filled dot: patient 1, filled square: patient 2 and filled triangle: patient 3. Results are analyzed by a two-way ANOVA test and followed by a Tukey’s multiple comparisons test. Plasma K^+^ is influenced by the patient’s status (p<0.05). Total excretion of GDF15 is not influenced by the patient’s status.

**Table 1.**
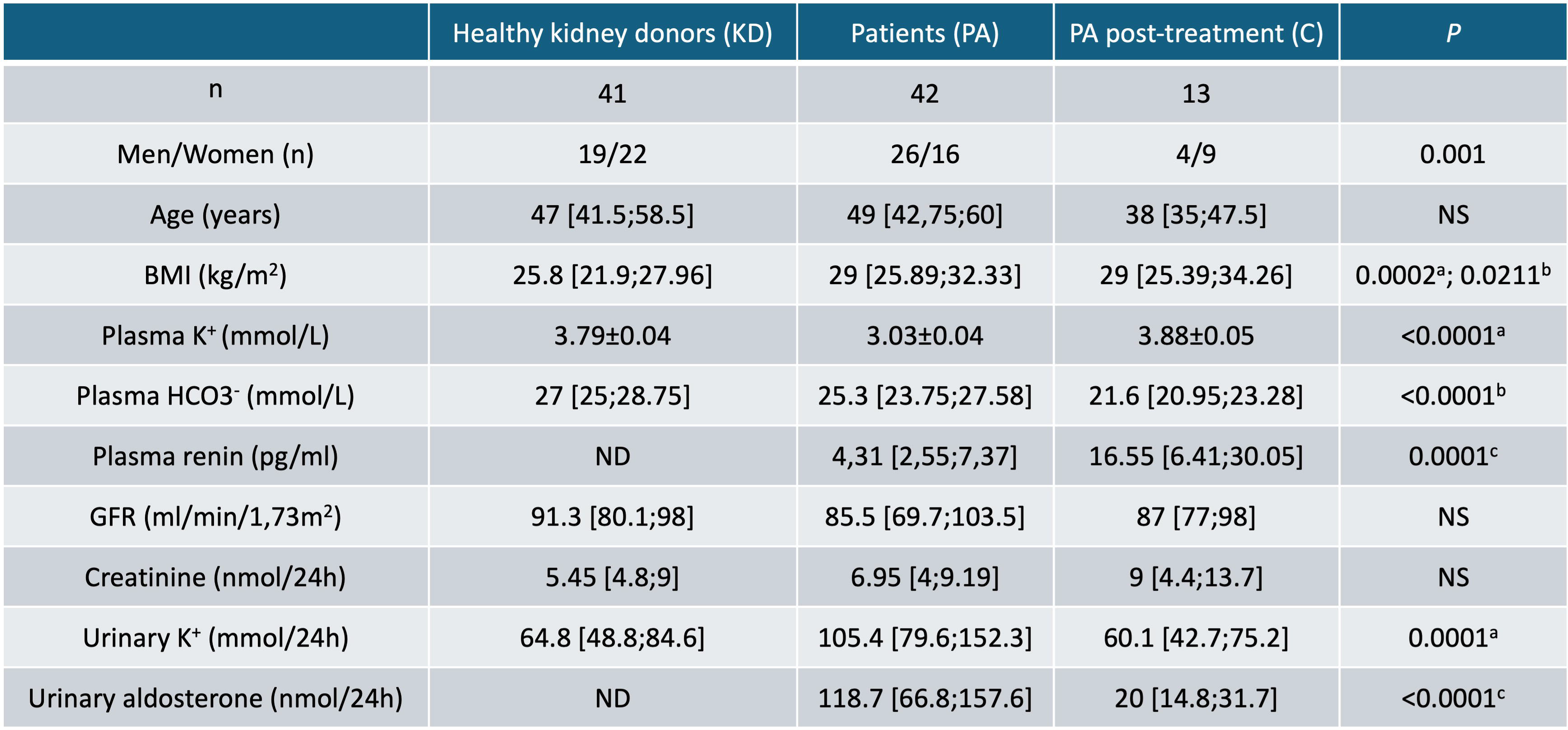
General characteristics of the cohort. Results are shown as mean ± s.e.m or median [Q1;Q3] according to the normality of the values. Comparisons between two groups were analyzed by a Mann-Whitney test. Comparisons between more than two groups were analyzed by a one-way ANOVA or its non-parametric equivalent, the Kruskal-Wallis test and followed by a Dunnett’s or Dunn’s multiple comparisons test. Qualitative variables were compared using the χ2 test. With ^a^KD vs PA, ^b^KD vs C, ^c^PA vs C.

To better investigate the relationship between GDF15 and clinical and biochemical parameters within PA patients, we added 39 patients with PA from a second center. These patients exhibit slightly higher mean plasma K^+^ values than in center 1 (3.25±0.04 vs 3.03±0.04, p<0.01, Supplemental data 1A) due to the inclusion of 13 normokalemic individuals. However, patients from both centers have a similar mean 24-hour urinary excretion and plasma GDF15 levels (Supplemental data 1B and C). Univariable analysis (Figure 2) revealed that 24-hour urinary GDF15 excretion is around 2-3-fold higher in men than in women (p<0,01) and is correlated with age (rho=0.36, p<0.01), BMI (rho=0.36, p<0.01), 24-hour urinary K^+^ (rho=0.33, p<0.01) and 24-hour urinary aldosterone (rho=0.35, p<0.01). Interestingly, GDF15 excretion is not correlated with plasma K^+^ values (rho=-0.16, p=0.18).

**Figure 2.**
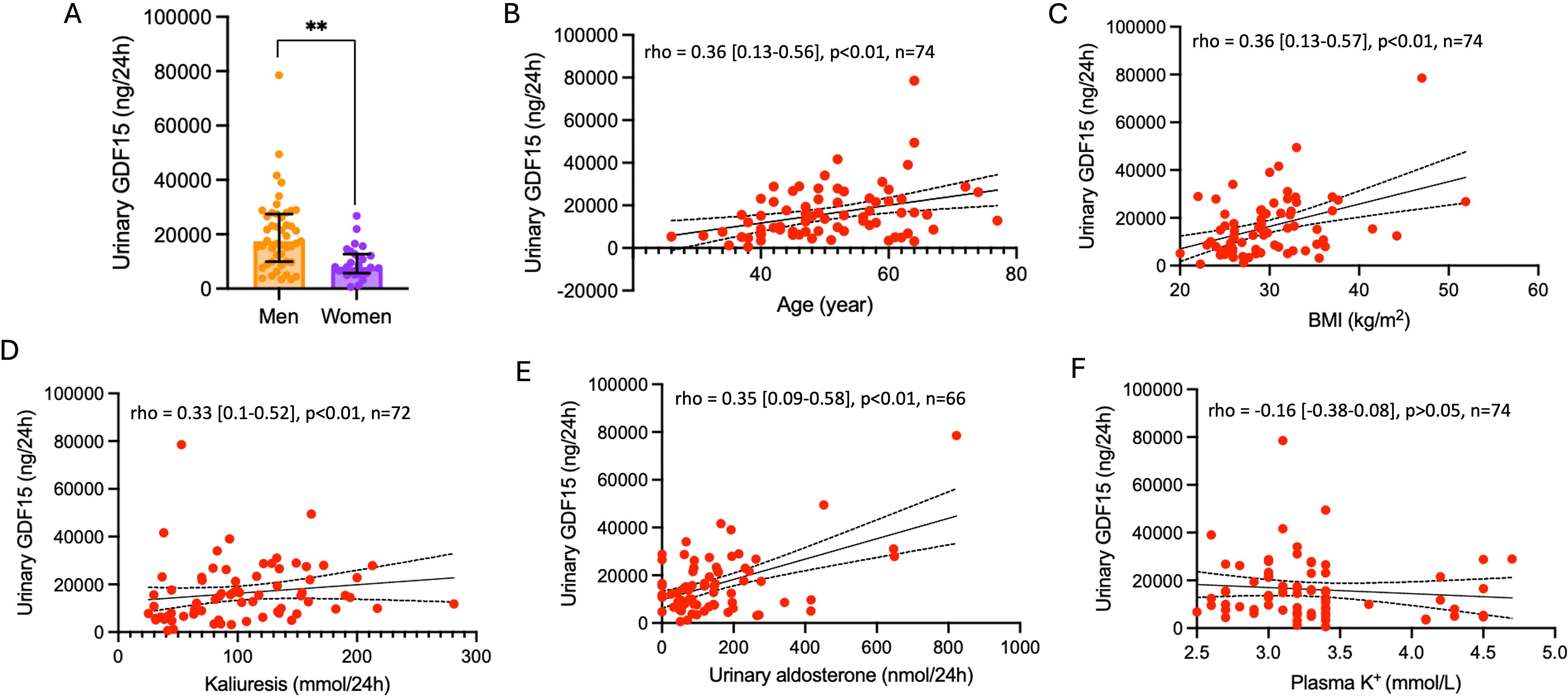
Urinary GDF15 as a function of sex, age, BMI, urinary K^+^ levels, urinary aldosterone and plasma K^+^ levels. A) Urinary GDF15 levels in men and women patients. Results are shown as median with interquartile range and analyzed by a Mann-Whitney test. **p<0.01. B-F) Continuous variables were compared using Spearman correlation. Each dot represents one patient.

To identify determinants of 24-hour urinary GDF15 levels, we used a hierarchical linear regression approach including sex, age, BMI, plasma K^+^ values, 24-hour urinary K^+^ and 24-hour urinary aldosterone as confounders and explanatory variables. We tested different models (Table 2) from the simplest (with only the sex as confounder) to the most complete one (including, sex, age, BMI, plasma K^+^ values, 24-hour urinary K^+^ and 24-hour urinary aldosterone). The Akaike Information Criterion (AIC) that decreased with the complexity of the models indicates that the most complete one is the best (F[6, 57]=10.73, p<0.01) and explains 38.3% of the variance of the urinary GDF15 in patients.

**Table 2.**
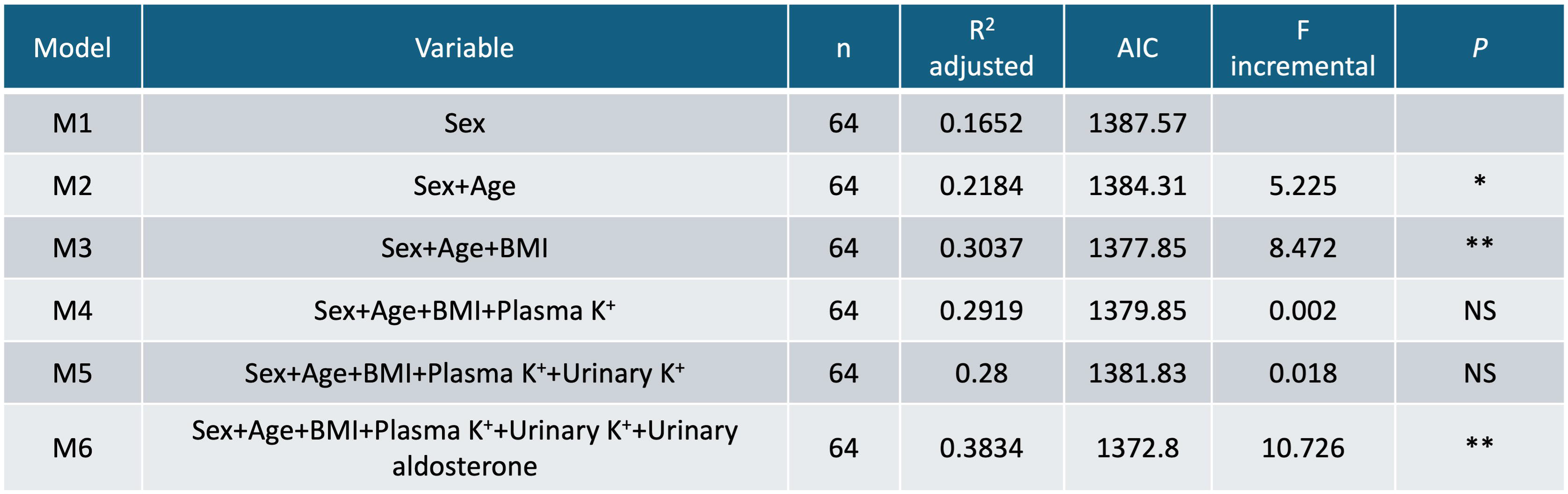
Independent factors associated with urinary GDF15 in patients with primary aldosteronism. The model is adjusted with sex, age, BMI, plasma K^+^, urinary K^+^ and urinary aldosterone. n the number of patients included, R^2^ adjusted = 0,3834 and AIC = 1372,8. *p<0.05 and ** p<0.01. Sex, BMI and urine aldosterone are the main determinant of urinary GDF15 in PA patients.

Among these variables, sex (ß=-6201.5 ng/24h, p<0.05), BMI (ß=501.4 ng/24h, p<0.05) and 24-hour urinary aldosterone (ß=32.18 ng/24h, p<0.01) were the main determinants of urinary GDF15 excretion contributing to 16.5%, 8.5% and 10.3%, respectively to its variance (Table 3).

**Table 3.**
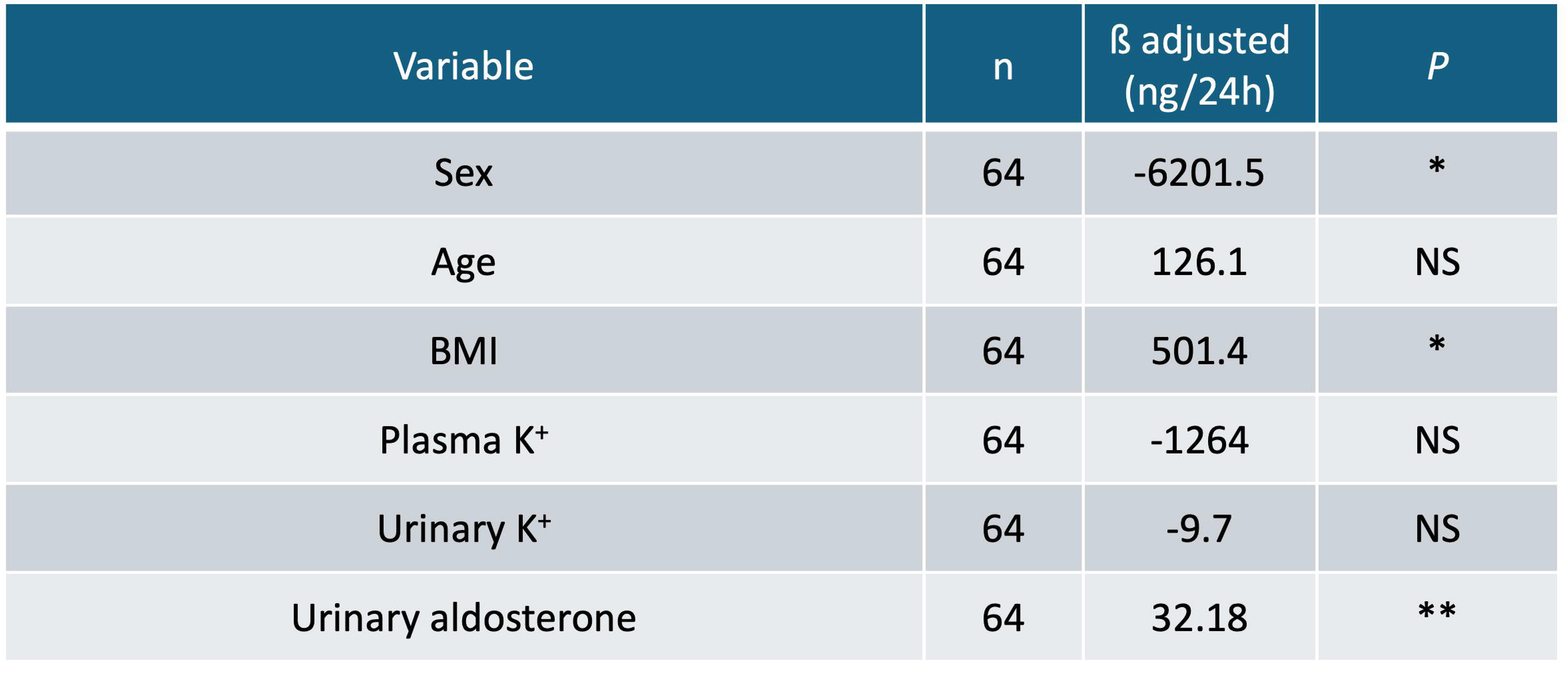
Determinants of the variance of urinary GDF15 in PA patients. A multiple linear regression was performed with sex, age, BMI, plasma K^+^ level, urinary K^+^ and urinary aldosterone as explanatory variables. Adjusted regression coefficients (ß) were calculated for each variable, representing the change in urinary GDF15 excretion for each one-unit increase in the respective predictor or the effect of being a woman. Patients with missing data were excluded from this analysis. For all analyses, differences with p<0.05 were considered significant.

### GDF15 excretion and expression are increased in PA murine model

To investigate the involvement of GDF15 in PA, we develop a mouse model of chronic mineralocorticoid excess as already described^24^. As shown in Figure 3A, the unique intramuscular injection of DOCP leads to a sustained increase of 11-deoxycorticosterone in plasma of treated animals for up to 28 days. To check the efficiency of the DOCP treatment, we measured the accumulation of fluid by TD-NMR analysis (Figure 3B) which is significantly increased by 14% after 1 day treatment. In parallel, the Na^+^ excretion significantly decreased by 18% (p<0.01, Figure 3C) and the K^+^ excretion significantly increased by 4.5% (p<0.05, Figure 3D) which leads to a significant decrease of the plasma K^+^ level compared to untreated animals (3.5±0.1 mM vs 4.3±0.1 mM, respectively, p<0.01, Figure 3E). Therefore, as expected, the treatment with DOCP induced an increase of extracellular volume, sodium retention, a loss of K^+^ and a marked decrease of plasma K^+^ values. The 24-hour urinary excretion of GDF15 is 2-fold increased after DOCP injection up to day 9 and then returns to normal values (Figure 4A). Renal expression of GDF15 is not modified at days 2 and 9 post-treatment (Figure 4B), however, when we investigate the cortical collecting ducts (CCD) or the outer-medulla collecting ducts (OMCD), we found a significant increase of GDF15 expression at day 2.

**Figure 3.**
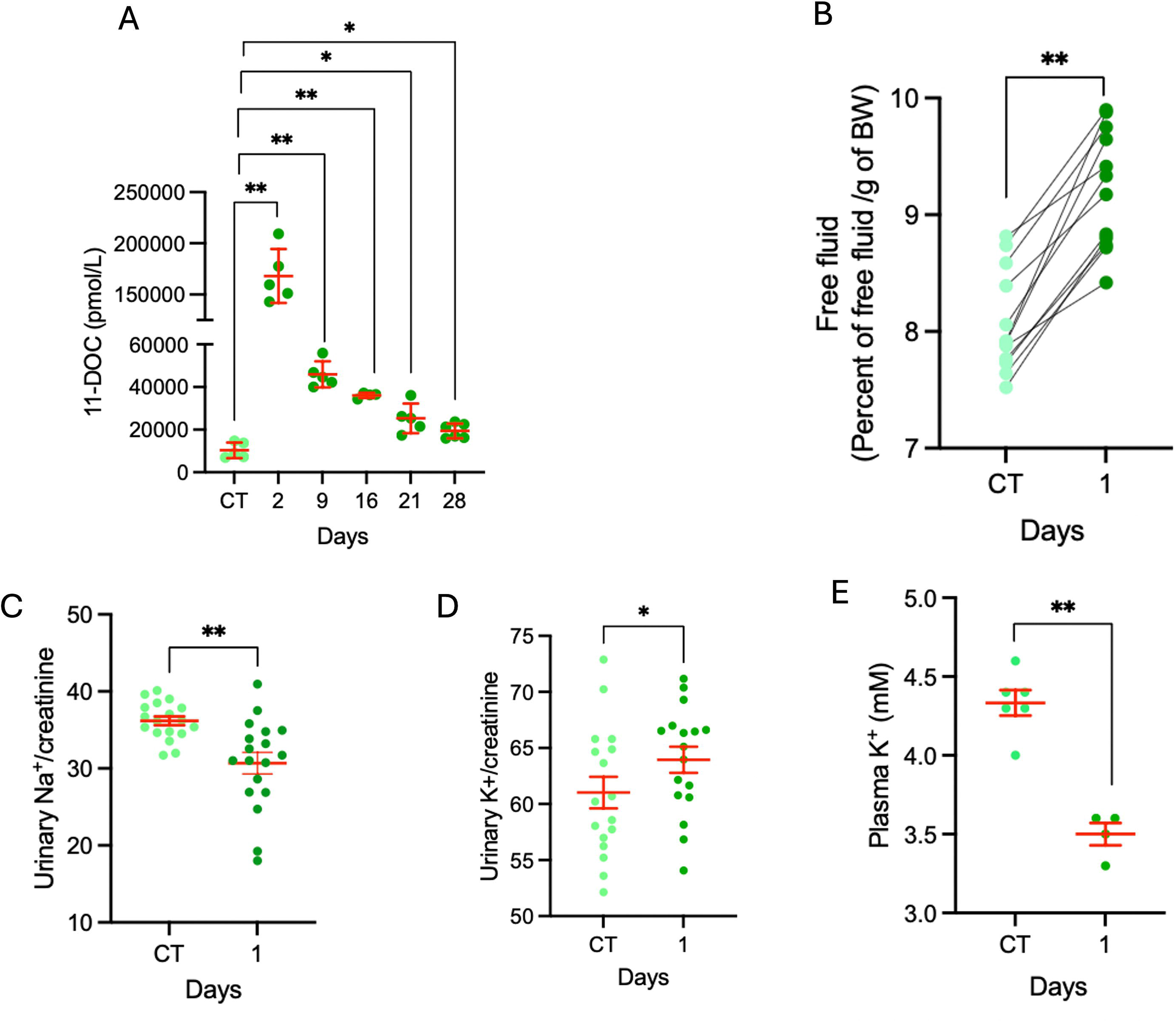
Validation of the murine model of primary aldosteronism. A) Plasma 11-desoxycorticosterone under control conditions and at day-2, 9, 16, 21 and 28 post treatment. Results are shown as mean ± s.e.m (n= 5-6) and analyzed by a Brown-Forsythe and Welch ANOVA test and followed by a Dunnett’s T3 multiple comparisons test with CT as control group. **p<0.01; *p<0.05. B) Percent of free fluid per gramme of body weight in WT mice under control condition and day-1 post treatment (n= 13) and analyzed by a paired Student t test. **p<0.01. C-D) Urinary Na^+^ and K^+^ normalized by urine creatinine in WT mice under control condition and day-1 post treatment. Results are shown as mean ± s.e.m (n= 17-18) and analyzed by a paired Student t test. **p0.01; *p<0.05. E) Plasma K^+^ values of WT mice under control conditions and day-1 post treatment. Results are shown as mean ± s.e.m (n= 4-6) and analyzed by an unpaired Student t test. **p<0.01.

**Figure 4.**
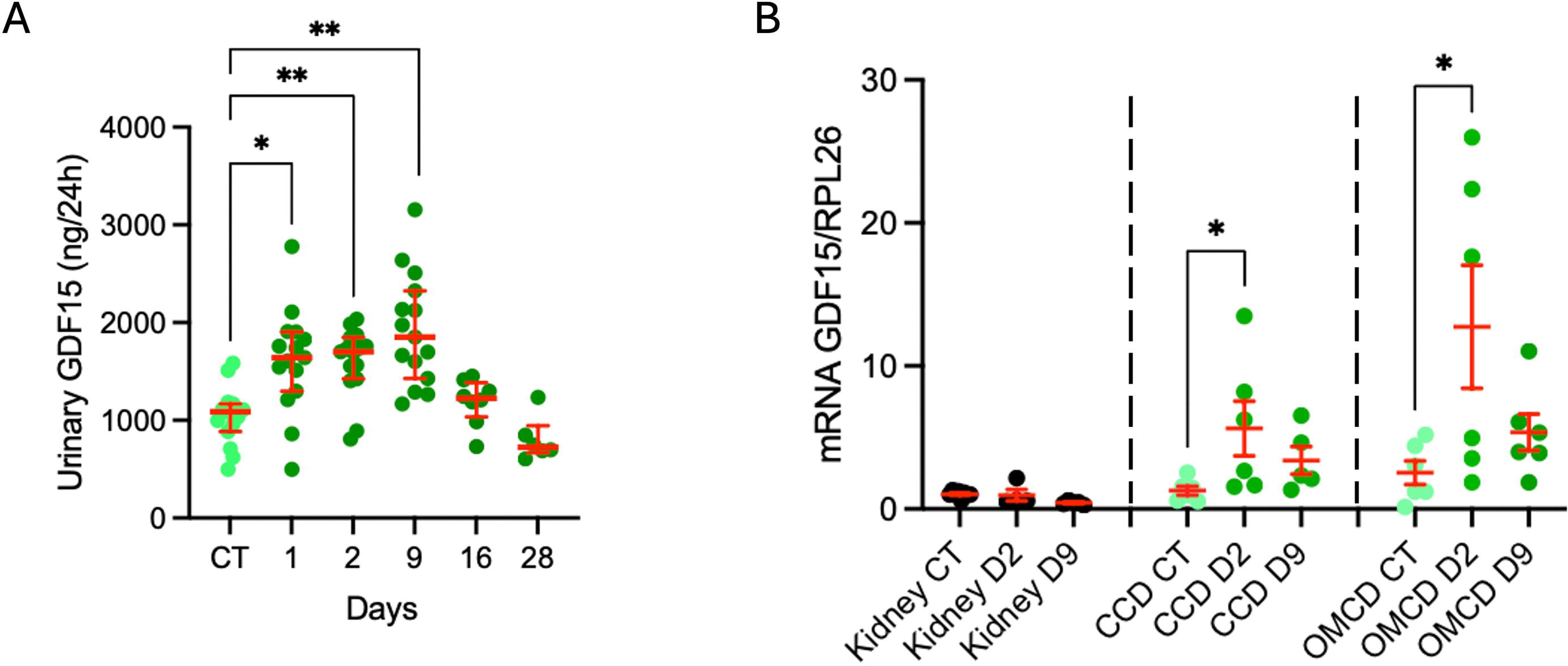
GDF15 is produced in response to DOCP injection. A) Total excretion of GDF15 under control conditions and day-1, 2, 9, 16 and 28 days post treatment. Results are shown as median with interquartile range (two series n= 6-15) and analyzed by a Kruskal-Wallis test and followed by a Dunn’s multiple comparisons test. **p<0.01; *p<0.05. B) mRNA expression of GDF15 in kidney, isolated cortical or outer medullary collecting duct (CCD and OMCD) of WT mice under control conditions, day-2 and 9 post treatment. Results are shown as mean ± s.e.m (n= 4-8) and analyzed by a one-way ANOVA test and followed by a Dunnett’s multiple comparisons test with CT as control group. *p<0.05.

### The absence of GDF15 modifies renal adaptation to mineralocorticoid excess

We measured the impact of chronic mineralocorticoid excess (day 16 post-treatment) by analyzing the global modification of the renal transcriptome in WT and GDF15KO mice. As shown in Figure 5A, the treatment significantly modifies the expression of 49 genes in WT mice (30 are upregulated and 19 are downregulated) based on a 2-fold cutoff. Upregulated genes in WT mice included *Bmal 1*, *Kcnj1* and *Pvalb* already known to be induced by chronic aldosterone excess whereas *Ren1*, for instance, is downregulated. Interestingly, when we analyzed specific GO gene sets representing the collecting duct cell population (Figure 5B), we found that those specific to ICA and PC of the medulla^25^ were significantly affected with most of their representative genes being upregulated (Figure 5C). In GDF15KO mice, the treatment significantly modifies the expression of 32 genes (27 are upregulated and 5 are downregulated) based on a 2-fold cutoff (Figure 5D). Analysis of the same GO gene sets revealed that the ICA and PC of the medulla were also significantly affected (Figure 5E) by the mineralocorticoid excess, but with a completely inverse effect, most of the genes were downregulated (Figure 5F).

**Figure 5.**
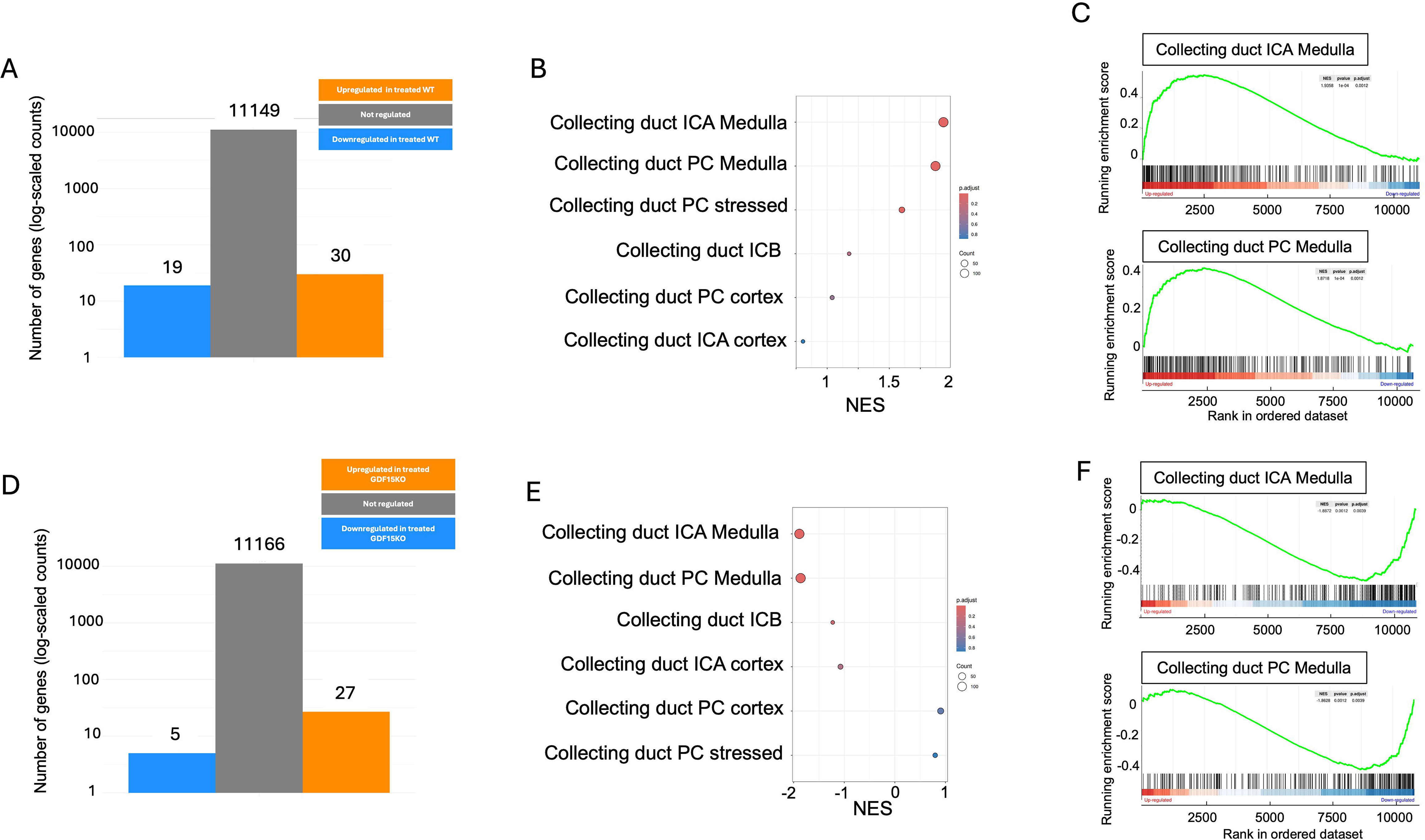
Differential expression analysis in response to DOCP injection both in WT and GDF15KO mice. A, D) Number of significantly up- and down-regulated genes respectively in WT and GDF15KO mice in response to DOCP treatment with log2FC=1 and p<0.05. B, E) GSEA using the C8 Cell type signature gene set collection from the molecular signatures data base (MSigDB) in WT and GDF15KO mice in response to DOCP treatment. The bubble size corresponds to the number of genes related to the set. The color intensity of the bubble corresponds to the p.value. C, F) Enrichment score plot of type A intercalated cells and principal cells of the medulla of the collecting duct in WT and GDF15KO mice in response to DOCP treatment.

### ICA number is increased by mineralocorticoid excess through a GDF15-dependent pathway

In K^+^ restriction, we demonstrated that GDF15 induced the increase number of ICA in OMCD^19^. Moreover, RNA-seq results indicate that chronic DOCP treatment has an inverse impact, in WT and GDF15KO mice on the transcriptomic characteristics of this cell type in the medulla. We therefore quantified the number of ICA in response to DOCP treatment in both WT and GDF15KO mice. For this purpose, we isolated the OMCD and performed an immunolabelling with an anti-anion exchanger 1 (AE1) antibody, a specific marker of ICA (Figure 6A) as described previously^18^. To measure the total number of cells we labelled the nuclei with DAPI. As shown in Figure 6B, the DOCP treatment did not modify the number of cells/mm either in WT or GDF15KO mice. However, the percentage of AE1+ cells in WT mice increased from 22.0±0.5% in the untreated group to 24.3±0.6% on day 16 (p<0.05) and to 25.7±0.8% on day 28 (p<0.01) post-treatment (Figure 6C). In GDF15KO mice, the percentage of ICA in the untreated group is similar to that in WT mice (22.5±0.6%) and never increased in response to DOCP treatment (Figure 6C). ICA are normally completely isolated in the epithelium, therefore, the observation of two adjacent ICA (a doublet) highlights a recent cell division. As shown in Figure 6D, the number of ICA doublets is significantly increased by the DOCP treatment in WT mice on day 16 compared to the control group (24±2 doublets/mm and 12.5±1.3 doublets/mm, respectively, p<0.01). In GDF15KO mice, the number of doublets is not modified by the treatment at any time (22.2±1.5 doublets/mm in untreated group and 22.6±2.3 doublets/mm at day 16 post-treatment). The stimulation of the HKA2 has been shown to be important for the proliferation of the ICA^19^. Here, we showed that the expression of HKA2 tends to increase on days 16 and 28 (p=0.07 at day 28) when the number of ICA also increased. However, it remained completely stable in GDF15KO mice during DOCP treatment (Figure 6E).

**Figure 6.**
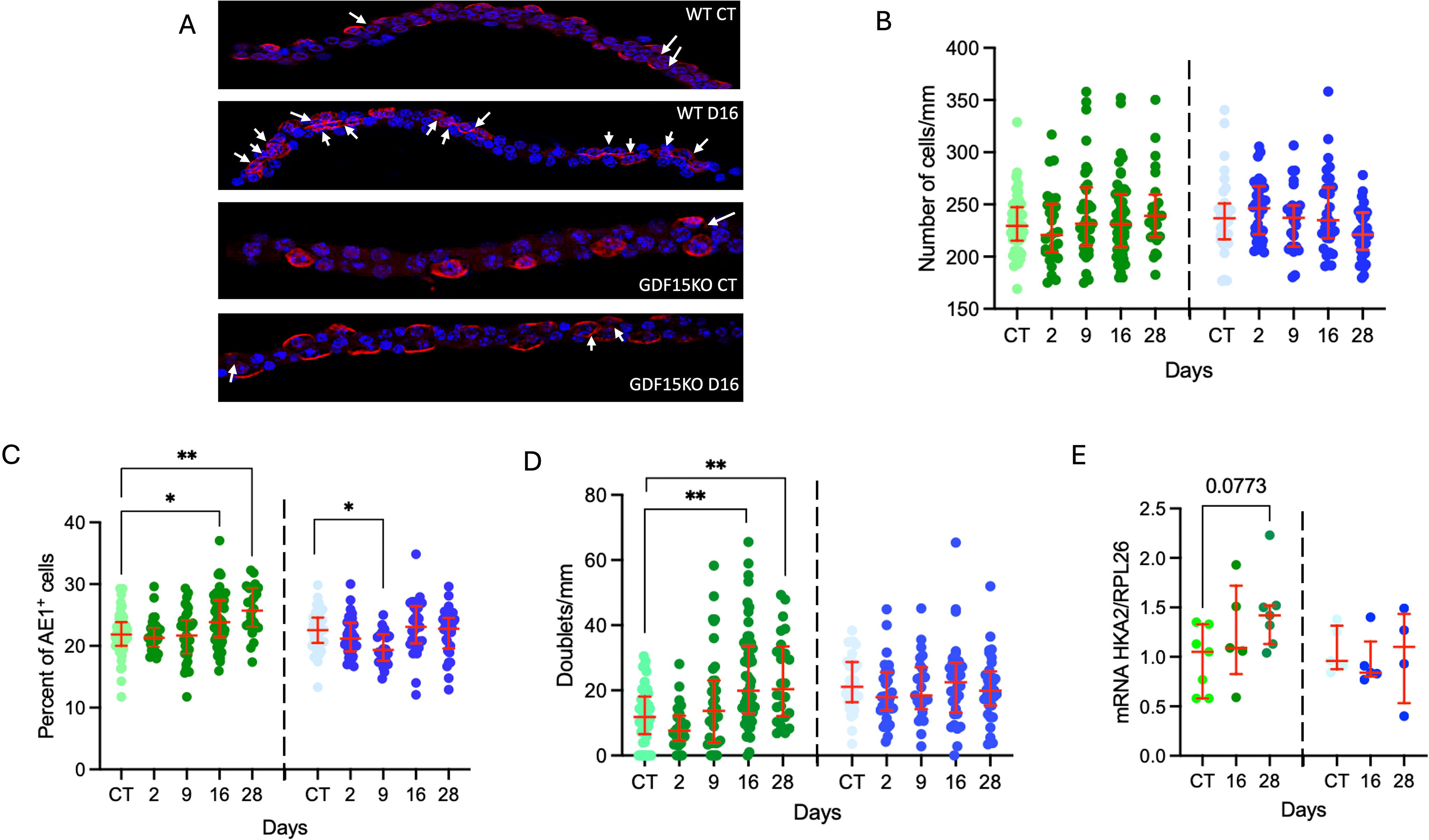
Plasticity of outer medullary collecting duct. A) Immunofluorescence pictures obtained by confocal microscopy from isolated OMCD from WT or GDF15KO mice under control conditions and day-16 post treatment, labeled with DAPI (blue) and anti-AE1 antibody (red). Stacks were used to reconstruct tubule in 3D and count the number of nucleus and doublets of type A intercalated cells. White narrows represent doublets of ICA. B) Number of cells in WT (green) and GDF15KO (blue) mice under control conditions and day-2, 9, 16, 28 post treatment. Results are shown as median with interquartile range (n= 24-55 tubules) and analyzed by a Kruskal-Wallis test or one-way ANOVA and followed by a Dunn’s or Tukey’s multiple comparisons test. No statistically significant differences were observed between groups p>0.05. C) Percent of AE1^+^ cells in WT (green) and GDF15KO (blue) mice under control conditions and day-2, 9, 16, 28 post treatment. Results are shown as median with interquartile range (n= 24-55 tubules) and analyzed by a Kruskal-Wallis test or one-way ANOVA and followed by a Dunn’s or Holm-Sidak’s multiple comparisons with CT as control group. **p<0.01; *p<0.05. D). Number of doublet (at least two AE1^+^ which cell-cell contact) in WT (green) and GDF15KO (blue) mice under control conditions and day-2, 9, 16, 28 days post treatment. Results are shown as median with interquartile range (n= 24-55 tubules) depending on normality and analyzed by a Kruskal-Wallis test and followed by a Dunn’s multiple comparisons test with CT as control group. **p<0.01. E) mRNA expression of HKA2 in kidney of WT (green) and GDF15KO (blue) mice under control conditions and day-16 and 28 post treatment. Results are shown as median with interquartile range (n= 4-7) and analyzed by a Kruskal-Wallis test and followed by a Dunn’s multiple comparisons test with CT as control group. No statistically significant differences were observed between groups p>0.05.

### The absence of GDF15 exacerbates hypokalemia in PA murine model

Since we observed that the number of ICA in OMCD did not increase following DOCP injection in GDF15KO mice, we investigated whether it affects the K^+^ homeostasis compared to WT animals. As shown in Table 4, WT and GDF15KO mice exhibit similar weight, food intake and urine volume before and during the treatment. As previously shown in Figure 3D, the 24-hour urinary K^+^ excretion is slightly but significantly increased in WT mice one day post treatment and returned to a similar level compared to the untreated mice by day 2 post-treatment. Similarly, GDF15KO mice did not exhibit any modification of their 24-hour urinary K^+^ excretion from days 2 to 28 post-treatment (Figure 7A). Maintaining potassium balance comes at the expense of plasma K^+^ levels and muscle K^+^ content. Indeed, as shown in Figure 7B, DOCP treatment induced a time- and genotype-dependent decrease of plasma K^+^ levels. Plasma K^+^ values are both impacted by duration of the treatment (p<0.01) and by the genotype (p<0.01) but not by their interaction. Thus, DOCP treatment induced a significant decrease in plasma K^+^ levels in WT mice that stabilized around 3.5 mM on days 2 and 9 post-treatment whereas the GDF15KO mice reached 3.05±0.07 mM on day 9. In WT mice, the plasma K^+^ level returned to values similar to those of the untreated group from days 16 to day 28 (around 4 mM) whereas in GDF15KO mice, it stagnated at 3.5 – 3.6 mM which is 1 mM below the GDF15KO control mice. Muscle K^+^ content is also affected by DOCP treatment depending on the duration of the treatment (p<0.01) and on the interaction between the post-treatment period and genotypes (p<0.01) (Figure 7C). In WT mice, muscle K^+^ content drops by 36% on day 9 but returns to control level by day 16, in parallel with the renal adaptation and the increase of ICA number. This may indicate that K^+^ storage is used to maintain plasma K^+^ values within an acceptable range before renal adaptation occurs. During this period (day 9), the muscle K^+^ content of GDF15KO mice remains unchanged and is significantly higher than that of WT mice (p<0.01) despite their lower plasma K^+^ level. GDF15KO mice eventually depleted their muscle K^+^ storage by day 16 indicating that a GDF15-independent and late-stage mechanism had been triggered to preserve their plasma K^+^ level. Noteworthy, neither WT nor GDF15KO mice modified their intestinal absorption of K^+^ during the DOCP treatment (Supplemental data 2A). However, the differential gene expression between WT and GDF15KO treated mice showed that *Kcnj1* (encoding for ROMK) is upregulated in WT (Supplemental data 2B). Indeed, DOCP inversely modulated *Kcnj1* expression in WT and GDF15KO mice (Supplemental data 2C). This suggest that GDF15KO mice may limit their K^+^ secretion by downregulating ROMK.

**Figure 7.**
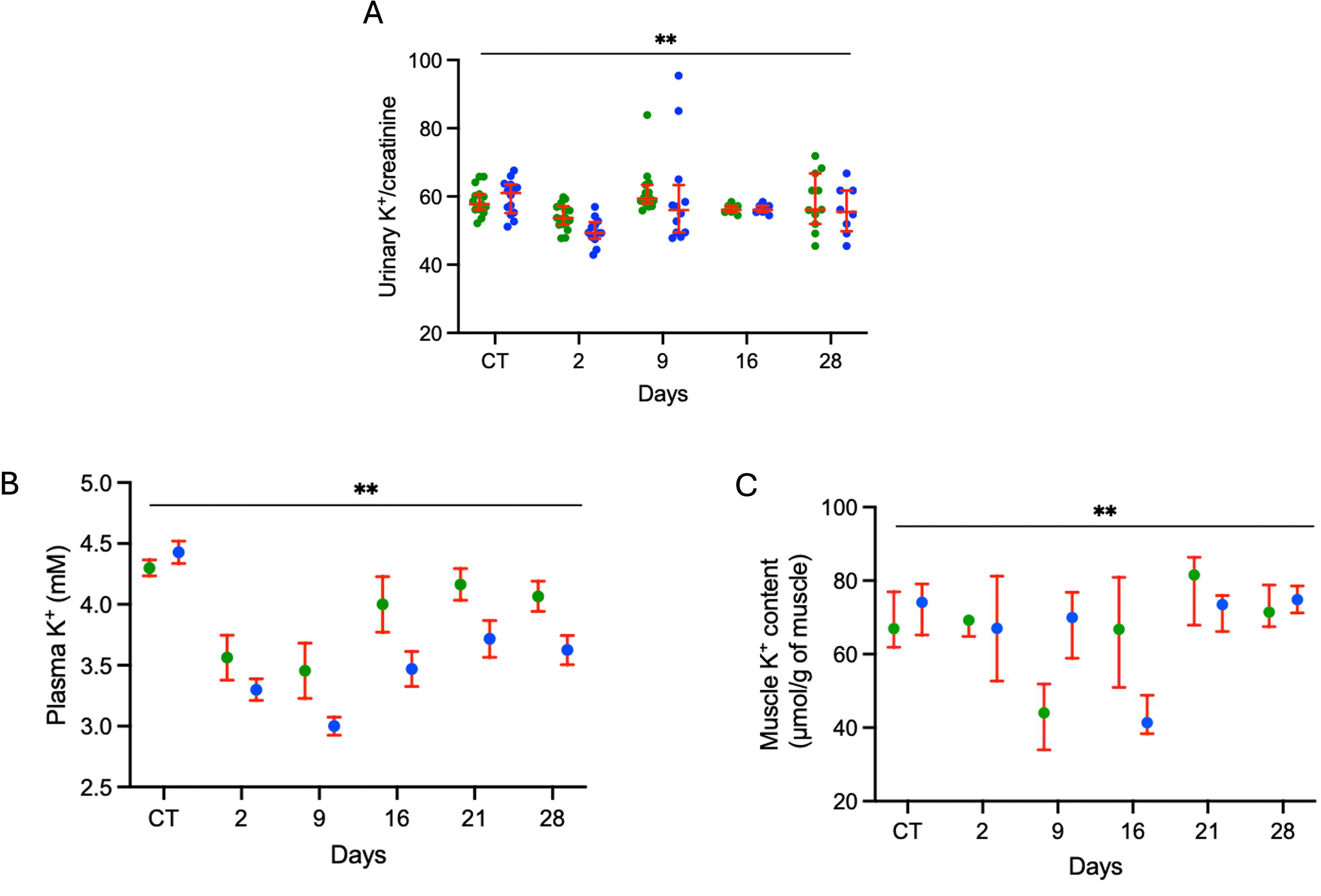
GDF15 is necessary to maintain plasma K^+^ in response to primary aldosteronism. A) Urinary K^+^ in WT (green) and GDF15KO (blue) mice under control conditions and day-2, 9, 16, 21 and 28 post treatment. Results are shown as median with interquartile range (two series, n=7-16) and analyzed by a two-way ANOVA and followed by a Sidak’s multiple comparisons test. Urinary K^+^ is significantly impacted by the treatment (p<0.01), not by the genotype and the interaction of both. B) Plasma K^+^ in WT (green) and GDF15KO (blue) mice under control conditions and day-2, 9, 16, 21 and 28 post treatment. Each symbol represents the mean of 6-12 mice. Results are shown as mean ± s.e.m (two series) and analyzed by a two-way ANOVA and followed by a Sidak’s multiple comparisons test. Plasma K^+^ is significantly impacted by the treatment and the genotype (p<0.01) by not by the interaction of both. C) Muscle potassium content in WT (green) and GDF15KO (blue) mice under control conditions and day-2, 9, 16, 21 and 28 post treatment. Each symbol represents the mean of 4-11 mice. Results are shown as median with interquartile range (two series) and analyzed by a two-way ANOVA and followed by a Sidak’s multiple comparisons test. Muscle K^+^ content is significantly impacted by the treatment (p<0.01), not by the genotype but by the interaction of both (p<0.01).

**Table 4.**
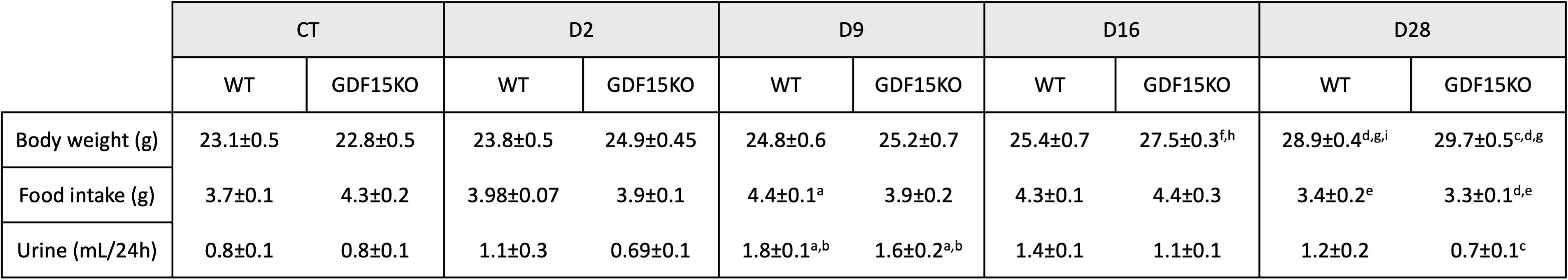
Physiological parameters of WT and GDF15KO mice before injection and day-2, 9, 16, 28 post treatment. Results are shown as mean± s.e.m (n= 7-10) and analyzed by a two-way ANOVA and followed by a Sidak’s multiple comparisons test. Body weight is significantly impacted by the genotype (p<0.01), the treatment (p<0.01) and the interaction of both (p<0.05). Food intake is significantly impacted by the treatment (p<0.01), not by the genotype and but by the interaction of both (p<0.05). Urine is significantly impacted by the genotype (p<0.01) and the treatment (p<0.01) not by the interaction. ^a^CT vs D9 ; ^b^D2 vs D9 ; ^c^D9 vs D28 ; ^d^CT vs D28 ; ^e^D16 vs 28 ; ^f^CT vs D16 ; ^g^D2 vs D28 ; ^h^D2 vs D16 and ^i^WT vs KO.

### Potassium influences GDF15 production in PA murine model

To test whether plasma K^+^ levels could determine the excretion of GDF15, we performed two types of experiment. First of all, mice were fed a high K^+^ diet (HK) one week before treatment with DOCP and were maintained under this diet for two days post-treatment. As shown in Figure 8A, as expected, mice under normal K^+^ diet significantly lowered their plasma K^+^ values (p<0.05) whereas those under HK diet maintained their plasma K^+^ values at level similar to the untreated mice (5.2±0.4mM vs 4.6±0.3 mM, respectively). The HK diet did not modify the GDF15 excretion before treatment but reduced the increase of GDF15 excretion by 26% 2 days post-treatment (Figure 8B). Secondly, to get closer to the clinical situation, we first treated the mice with DOCP to induce PA. On day 2 post treatment, some of mice were fed with HK diet until day 9. As previously shown in Figure 7B and as expected, DOCP treatment significantly decreased plasma K^+^ values only in mice fed a control diet whereas those fed a HK diet remained normokalemic. As already shown (Figure 4A), GDF15 excretion is 1.6-fold increased (p<0.01), 9 days post treatment compared to the untreated group. However, mice fed a HK diet did not increase their GDF15 excretion compared to the untreated group (Figure 8D). Altogether, these results indicate that plasma K^+^ levels and/or the K^+^ intake modulate GDF15 production in response to mineralocorticoid excess.

**Figure 8.**
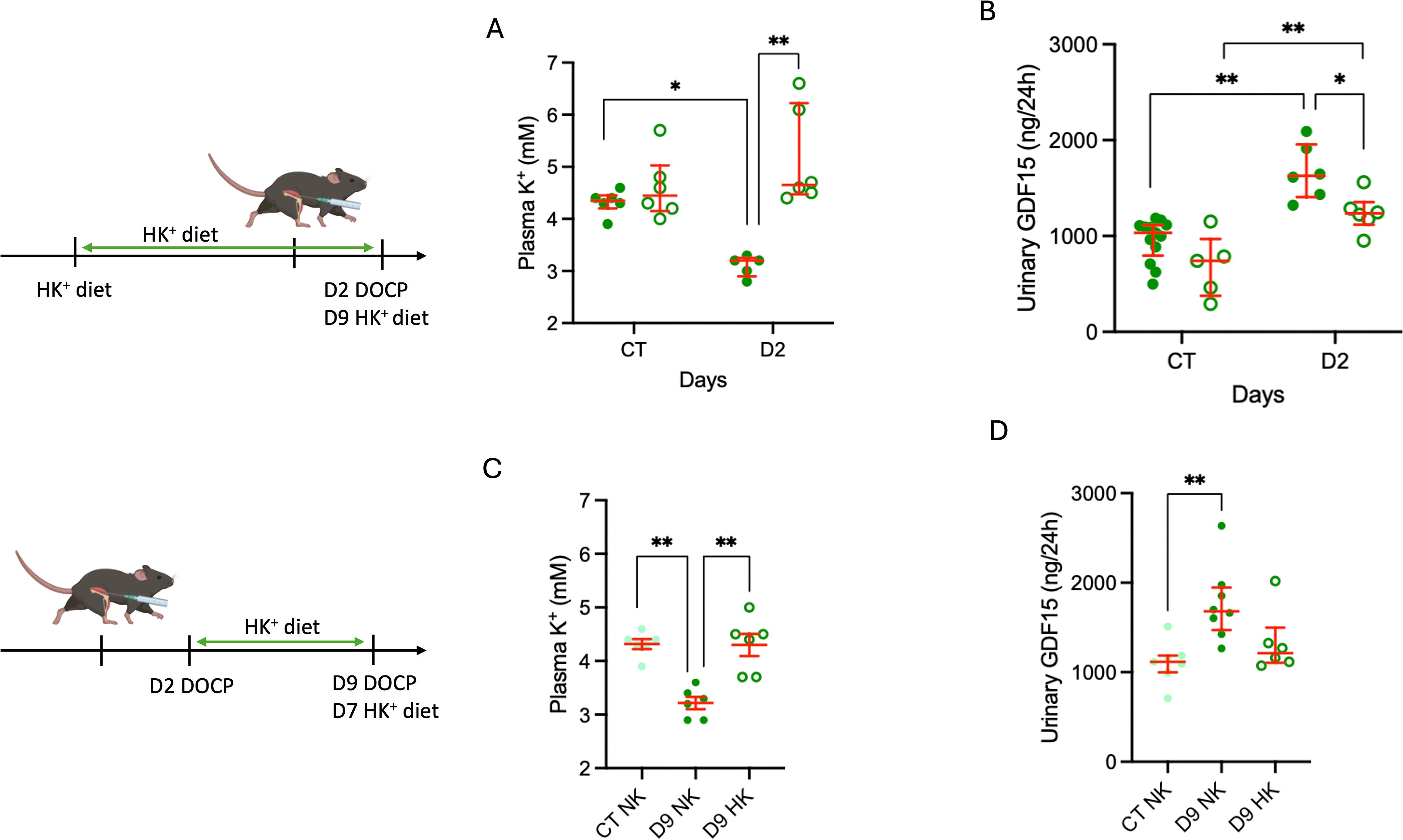
Potassium influences the production of GDF15 in primary aldosteronism murine model. A) Plasma K^+^ in control WT mice under control (filled symbols) or high K^+^ diet (open symbols) for one week and at day-2 post treatment. Results are shown as median with interquartile range (n=5-6) and analyzed by a two-way ANOVA and followed by a Tukey’s multiple comparisons test. **p<0.01; *p<0.05. B) Urinary GDF15 in control WT mice under control (filled symbols) or high K^+^ diet (open symbols) for one week and at day-2 post treatment. Results are shown as median with interquartile range (n=6-10) and analyzed by a two-way ANOVA and followed by a Tukey’s multiple comparisons test. **p<0.01; *p<0.05. C) Plasma K^+^ in WT mice in control condition and at day-9 post treatment under control diet (filled symbols) or high K^+^ diet since day-2 of DOCP treatment (open symbols). Results are shown as mean±s.e.m (n=6) and analyzed by a one-way ANOVA and followed by a Tukey’s multiple comparisons test. **p<0.01. D) Urinary GDF15 in WT mice in control and at day-9 post treatment under control diet (filled symbols) or high K^+^ diet since day-2 of DOPC treatment (open symbols). Results are shown as median with interquartile range (n=6-10) and analyzed by a Kruskal-Wallis test and followed by a Dunn’s multiple comparisons test with CT NK as control group. **p<0.01.

## Discussion

In this study, we investigated whether the growth factor GDF15 may be involved in the regulation of plasma K^+^ levels in the context of primary aldosteronism both in patients and in a pharmacological mouse model. Notably, our study has limitations, including the relatively small number of PA patients, particularly in the post-treatment group (n=13), and the initial cohort (center 1) was enriched in hypokalemic PA patients to focus on cases with the most severe hypokalemia which limited our ability to test multiple variables as determinants of the variance of urinary GDF15 levels. In primary aldosteronism (PA), plasma potassium levels are primarily determined by plasma aldosterone concentration, with a significant inverse correlation observed in PA^26^. Additionally, clinical and biological factors such as age, sex and systolic blood pressure are also correlated with the development of hypokalemia^27,28^. However, these factors do not fully explain the variability of the plasma K^+^ levels observed in these patients. In the current study, we demonstrated that GDF15, a growth factor identified for being involved in the reabsorption of K^+^^19^, was increased both in urine and plasma of PA patients. Pharmacologically-induced PA in mice showed that GDF15 is important for maintaining K^+^ homeostasis since its absence led to a more severe hypokalemia and a delayed adaptation to mineralocorticoid excess due to an inability to activate K^+^ reabsorption in an ICA-dependent manner. Our data suggest that GDF15 may help limit the development of hypokalemia in PA patients.

Beyond its role in PA, GDF15 has been implicated in a variety of physiological and pathological conditions. GDF15 is typically present at a low-levels in plasma, urine or tissues in normal or non-pathological conditions. However, its levels increase in many different circumstances ranging from pregnancy^29^ to physical exercise^30^, obesity/metabolic syndrome (for review^31^) and cancer (related to cachexia and resistance to treatment^32,33^) for the main examples. Regarding renal pathologies, urinary and plasma GDF15 levels were found increased in chronic and acute kidney disease and associated with comorbidities and pathological features^34,35^. Investigations using mouse models of CKD and AKI indicate that GDF15 may protect against fibrosis and may has a nephroprotective action through Klotho-dependent processes^36^. Moreover, plasma GDF15 level is negatively correlated with eGFR in healthy^37^ and CKD patients^38^. This is particularly relevant, as our PA patients have similar eGFR than the control group. This indicates that they had not developed kidney disease and, thus, the observed GDF15 increase was not related to renal dysfunction. Furthermore, BMI differences between groups cannot explain the GDF15 variations, since the PA post-treatment group—despite having a higher BMI than controls—showed similar GDF15 levels. Among the general characteristics known to influence plasma GDF15 level, the age and the sex have been shown to be of importance^39^, with higher levels in older individuals and men. In our study, the age is similar between the different groups, roughly in between 40 to 50 years old. The sex ratio differs between groups with a higher proportion of men in the PA patient group. However, in this age range, the difference in GDF15 levels between men and women is reported to be around 10%^39^. This cannot account for the increased observed here.

In our patients, 24-hour urinary aldosterone was a determinant of the variance of the levels of 24-hour urinary GDF15 which may suggest that GDF15 is an aldosterone-induced gene. However, we have shown that GDF15 is also overexpressed in conditions where plasma K^+^ is decreased but aldosterone levels are low, such as in dietary K^+^ restriction^19^. Furthermore, GDF15 did not appear to be a regulated gene in different studies where cultured PC models or in animal models were treated with aldosterone^40,41^. Therefore, the link between aldosterone and GDF15 is probably not direct and may be secondary to the development of hypokalemia at a very early stage. This is supported by our results showing that increasing K^+^ intake in the PA mouse model to prevent hypokalemia reduced or abolished the DOCP-induced GDF15 increase. Moreover, in WT DOCP-treated mice, urinary GDF15 excretion returns to a normal value when the plasma K^+^ value is restored although mineralocorticoid levels remained is still 4-times higher than in untreated mice.

These murine data complement our clinical findings, suggesting that GDF15’s role in K homeostasis is conserved across species. Interestingly, the GDF15KO mice partially succeeded in overcoming the excess of mineralocorticoid and returned to a plasma K^+^ value around 3.5 mM which, however, remained around 1 mM lower than that of the control group. This was not due to an increase in food intake (and therefore K intake) or to improved intestinal assimilation. We observed that during the late phase of the DOCP-treatment, muscle K^+^ content decreased in GDF15KO mice, which suggest that it may contribute to increase plasma K^+^ value. Analysis of differential gene expression between treated WT and GDF15KO mice showed that *Kcnj1* expression is inversely regulated between WT mice and GDF15KO mice being upregulated in WT mice and downregulated in GDF15KO mice. This may suggest that, when the GDF15-dependent activation of the K^+^ reabsorption pathway is lacking, there are mechanisms allowing a reduction of K^+^ secretion. However, it is obvious that reducing only K^+^ secretion is not sufficient. The increased number of ICA, at the detriment of the number of PC, is much more efficient since the ability for K^+^ secretion is de facto reduced while the ability for reabsorption is strongly stimulated.

Finally, our study identifies GDF15 as an active component of the potassium balance in the context of an excess of mineralocorticoid in patients and we demonstrate that its effects mediate the increased ability of the kidney to retain potassium limiting hypokalemia. Our data highlight the underappreciated role of ICA in K homeostasis. While PC are classically viewed as the primary regulators of K secretion, our findings suggest that ICA-mediated K reabsorption, stimulated by GDF15, represents a critical counterbalance to aldosterone-induced kaliuresis. Future studies should investigate whether GDF15 levels correlate with the risk of cardiovascular complications or kidney dysfunctions in PA.

## Acknowledgements

Physiological studies have been performed with the help of Gaëlle Brideau and Nadia Frachon from the “plateforme d’exploration fonctionnelle du petit animal” of the team “Physiologie Rénale and Tubulopathies” at the Centre de Recherche des Cordeliers. We are grateful for the technical assistance of the Centre d’Exploration Fonctionelle crews in the management of our colonies of mice. This study was supported by the Agence National de la Recherche (ANR) project ANR-21-CE14-0040-01.

---

**Supplemental data 1. The collecting center has no impact on GDF15 measurements.**

A) Plasma K^+^ in PA patients from center 1 and center 2. Results are shown as median with interquartile range and analyzed by a Mann-Whitney test. **p<0,01. B) Total excretion of GDF15 and C) plasma GDF15 in PA patients from center 1 or center 2. Results are shown as median with interquartile range and analyzed by a Mann-Whitney test. No statistically significant differences were observed between groups p>0.05.

**Supplemental data 2. Late adaptation to DOCP-treatment in GDF15KO mice.**

A) Stool potassium excretion in WT and GDF15KO mice in control conditions and at day-21 post treatment. Results are shown as mean±s.e.m (n=6-9) and analyzed by a two-way ANOVA and followed by a Tukey’s multiple comparisons test. No statistically significant differences were observed between groups p>0.05. B) Volcano plot of differential expression. Colored dots show genes with adjusted p<0.05. Blue dots represent down-regulated and orange dots represent up-regulated genes in WT treated mice compared to GDF15KO mice. C) Expression of *Kncj1* (count per million) in WT and GDF15KO mice in control conditions and at day-16 post treatment. n=5 and **p<0.01.

