## Supplementary figures and images for "Renal adaptation to primary aldosteronism requires GDF15 to control plasma potassium level in mice and human"

### Supplemental Fig 1

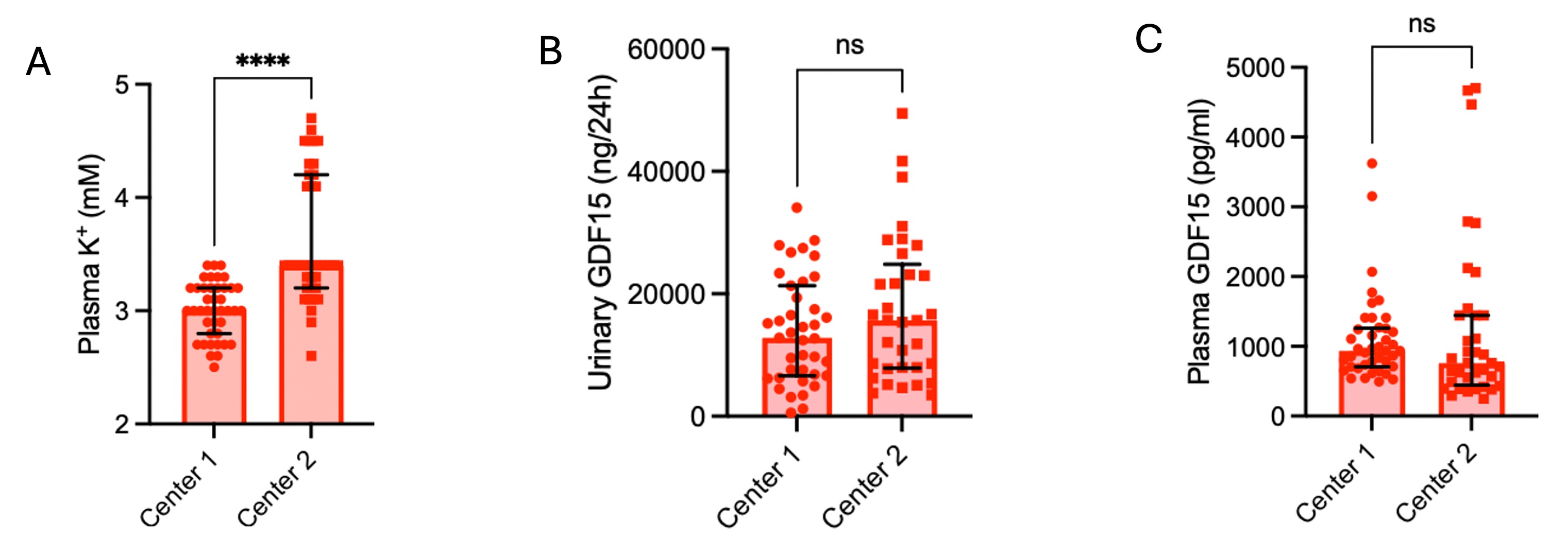

### Supplemental Fig 2

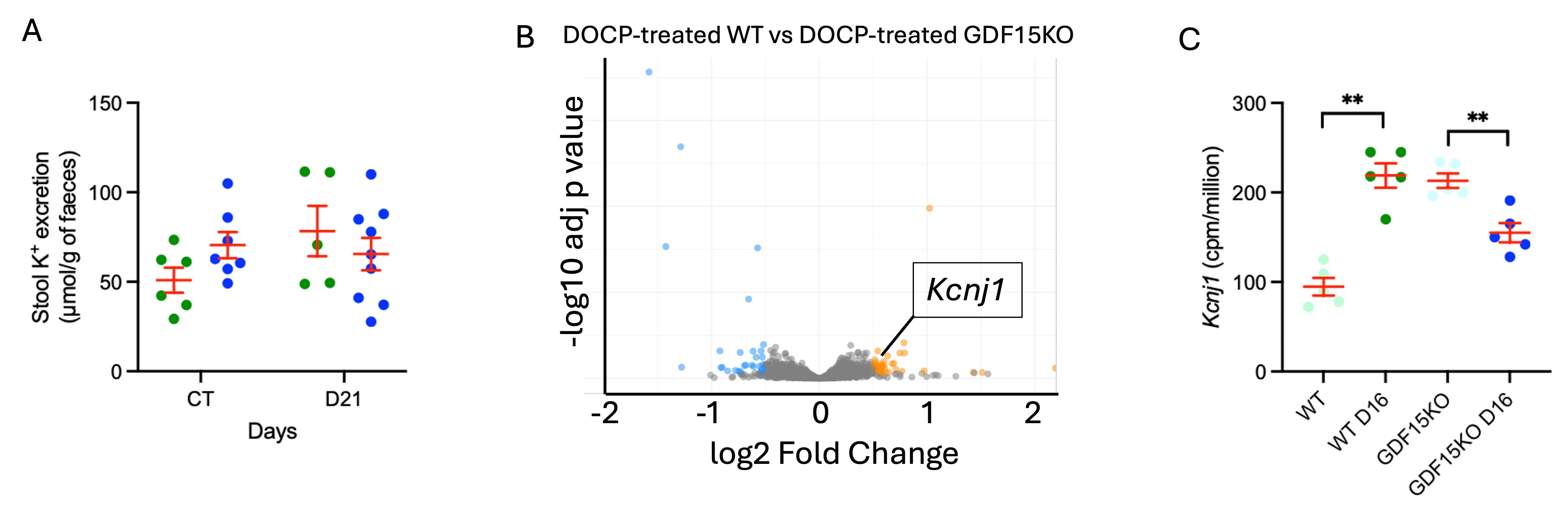
